# Seizure-associated spreading depression is a major feature of ictal events in two animal models of chronic epilepsy

**DOI:** 10.1101/455519

**Authors:** Fatemeh Bahari, Paddy Ssentongo, Jiayang Liu, John Kimbugwe, Carlos Curay, Steven J. Schiff, Bruce J. Gluckman

## Abstract

Spreading depression is characterized by slow, propagating wave of cellular depolarization (SD) and is wildly associated with migraine, stroke, and traumatic brain injury. Seizures and spreading depression (or spreading depolarization, SD) have long been reported to coincide in acute seizure induction experiments. However, SD has not been observed associated with spotaneous seizures in animal or clinical recordings. Recently, advances in acquisition systems for neurointensive care units have made routine observations of SD possible. In clinical epilepsy, SD has been suggested as a candidate mechanism for migraine/headache like events following seizures as well as for post-ictal generalized suppression. In animal models of epilepsy, seizure-induced brainstem SD has also been demonstrated as a mechanism of sudden unexplained death in epilepsy (SUDEP). The interplay between seizures and SD has also been suggested in computational models, where the two are components of the repetoir of neuronal activity.

However, the spatiotemporal dynamics of SD with respect to spontaneous seizures in chronically epileptic brain remains ambigous. We analyzed continuous long-term DC sensitive EEG measurements from two fundamentally different animal models of chronic epilepsy. We found that SD was associated with approximately one-third of all spontaneous seizures in each model. Additionally, SDs participated in the organization of seizure clusters. These findings demonstrate that the underlying dynamic of epileptic events is broader than seizures alone.

**Significance Statement:** Spreading depression is characterized by slow, propagating wave of cellular spreading depolarization (SD) and is wildly associated with migraine, stroke, and traumatic brain injury. Although recently the linkage between SD and induced seizures has been recognized, the mechanistic relationship between SD and spontaneous seizures remains poorly understood. Here, we utilized long-term, stable, near-DC measurements of the brain activity in two fundamentally different animal models of epilepsy to investigate the SD-seizure interplay. We found that SD is a frequent phenomenon in the epileptic brain, in these models is associated with more than a third of all seizures, and appears to connect seizures in seizure clusters. Although in one model SD stereotypically propagates out from a single focus in the hippocampus, depression of the field-potentials is observed synchronously across much of the hippocampus. These observations highlight the value of stable DC measurements for accurate understanding of SD and its propagation. We found that spontaneous ictal events that include both seizures and SD are frequent in animal models of epilepsy. These findings suggest that SD could be a valuable target for treatment and control of epilepsy.

## Introduction

Epileptic seizures were first associated with intense electrical activity measured from the brain in 1933 (Berger, 1933). Soon after, during attempts at acute experimental seizure induction, Leao discovered spreading depression of brain activity (Leao, 1944). The physiological underpinning of spreading depression is a spreading depolarization (SD) of brain cells that silences spontaneous or evoked activity. Since its discovery, SD has been shown to play a crucial role in multiple human pathological conditions including migraine, stroke, and traumatic brain injury (Lauritzen, 1994; Lauritzen et al., 2011; Hartings, 2017; Dreier et al., 2018).

Within epilepsy research, the study of SD has been mostly focused on seizure induction (Leao, 1944; Van Harreveld and Stamm, 1953; Vinogradova et al., 2005, 2006; Vinogradova, 2015; Khoshkhoo et al., 2017), genetic models (Aiba and Noebels, 2015; Loonen et al., 2019), and slice electrophysiology experiments (Fertziger and Ranck, 1970; Mody et al., 1987; Avoli et al., 1991; Köhling et al., 2003; Petzold et al., 2005; Olsson et al., 2006; Somjen et al., 2009; Maslarova et al., 2011; Eickhoff et al., 2014; Kramer et al., 2017). In vivo animal studies such as (Aiba and Noebels, 2015) and (Loonen et al., 2019) have brought to light the potential role of SD in sudden unexplained death in epilepsy (SUDEP).

Computational efforts have attempted to model the link between seizures and SD as well as its underlying mechanisms and suggested that the two are different components of the repertoire of neuronal membrane activity (Kager et al., 2000; Kager, 2002; Somjen, 2004; Wei et al., 2014a; Wei, et al., 2014b). However, the prevalence of and relationship between SD and spontaneous recurrent seizures remains poorly characterized and widely underappreciated.

The cellular electrical depolarization during SD is characterized with a large (15-35 mV), slow, and prolonged (10-30 s) negative shift in tissue potential that is observable in its true form only with sufficiently low frequencies (low-frequency cutoff of much less that f = 1/30 Hz). Stable near-DC measurements require sufficiently non-polarizing electrodes with impedance that is small relative to the input impedance and input bias current of the amplifier. Otherwise, the coupling between amplifier input and electrode-tissue interface forms an inherently nonlinear high-pass filter that significantly attenuates and distorts the low-frequency signatures of SD and makes them impossible to recover from the stop band. What is left are observations of activity suppression similar to clinical reports of post-ictal generalized suppression (Surges et al., 2011; Poh et al., 2012; Kim et al., 2015; Bateman et al., 2019; Marchi et al., 2019) or studies of mid-ictal and post-ictal suppression in animals (Smith et al., 2018). These observations might in fact be reflections of spreading events of cellular depolarization.

Recent improvement in clinical technologies for neurointensive care units (neuroICU) have enabled observations of SD in acute clinical measurements in various brain injury phenomena (Fabricius et al., 2008; Dreier et al., 2012, 2018). However, these improvements have not yet been implemented in either long-term clinical or experimental epilepsy recordings.

To address the relationship between spontaneous seizures and SD in chronic epilepsy, we examined SD occurrence and propagation in freely behaving epileptic animals. We utilized continuous long-term wide-band recordings sensitive to both seizures and SD dynamics (Ssentongo et al., 2017) to demonstrate that well-spaced spontaneous seizures are frequently associated with SD, and that seizures within clusters may be connected through SD. Our findings suggest that an epileptic event or ictus comprise combinations of dynamics of both seizure and SD.

## Materials and Methods

All animal work was approved by and performed under the oversight of the Institutional Animal Care and Use Committee at the Pennsylvania State University.

### Study Design

The relationship between spontaneous seizures and SD was investigated in two separate models of acquired epilepsy: first, retrospectively from recordings used to establish a murine model of post-cerebral malaria (post-CM) epilepsy (Ssentongo et al., 2017); and second, prospectively in the rat tetanus toxin (TeTX) model of temporal lobe epilepsy, in which we specifically implanted electrodes to resolve the spatial extent and evolution of these dynamics.

No explicit blinding was done with any of this data. Rather, signal-processing algorithms designed to independently detect seizure and SD events were automatically applied to all recordings regardless of cohort to provide unbiased results.

### Animal Surgery and Care

#### Post-Cerebral Malaria Epilepsy Model

All the data from the post-CM mice utilized here were acquired during development of a murine model of post-CM epilepsy previously described in (Ssentongo et al., 2017). Combinations of mouse and parasite strains were examined: male Swiss Webster (SW), C57BL/6 (Charles Rivers Laboratory), and CBA/CaJ (CBA, Jackson Laboratory) mice and both Plasmodium berghei ANKA (PbANKA) and Plasmodium berghei NK65 (Pb-NK65) parasites. Details of the experiments are included in (Ssentongo et al., 2017). Critical elements specific to the work presented here follow:

Animals were implanted with electrodes to monitor hippocampal, cortical, and nuchal muscle activity at least five days post-treatment following procedures described in (Ssentongo et al., 2017). A cohort of the animals also received electrocardiography (ECG) electrodes to monitor heart activity. Four stainless steel screw electrodes (#000 self-tapping, Morris Co.) were placed over frontal and S1 cortices to monitor cortical activity. For hippocampal local field potentials (LFPs), two custom-made 50 μm (diameter) ultra-low impedance micro-reaction chamber (μRC) electrodes were further enhanced by electrodeposition of iridium oxide films (EIROF) (Shanmugasundaram and Gluckman, 2017) to provide DC stability and high charge passing capacity. The μRC electrode provides a larger electrochemical interface between the electrode surface and tissue without increasing the electrode size. All electrodes were referenced to a stainless steel screw implanted at (AP −3.8, ML +3.5 – bregma referenced) and recordings were performed referentially. At the completion of the surgery, animals were returned to their individual home-cages with free access to food and water, and, following recovery, were cabled for continuous video and EEG monitoring.

#### Rat Tetanus Toxin Model of Temporal Lobe Epilepsy

Male and female Long-Evans rats, weighing 250-350 grams, were implanted with recording electrodes and received tetanus toxin injections during the same surgical procedure following methods described previously (Sedigh-Sarvestani et al., 2014). Critical elements specific to the work presented here follow: To induce epilepsy, 10-13 nano-grams of tetanus toxin (Santa Cruz Biotechnology, CAS 676570-37-9) dissolved in 1.3 microliters phosphate buffered saline (PBS) mixed with 2% bovine serum albumin (BSA) were injected into the left ventral hippocampus (AP −5.15, ML +5.35, DV −7.65 mm) through a 30-gauge flexible cannula over 15 minutes.

Recording electrodes included stainless steel screws for reference and ground at coordinates of (AP −6.5, ML ±4), and for measurements of electrocorticogram (ECoG) at coordinates of (AP +1.5, ML ±4 mm) and (AP −2, ML ±3 mm). ECoG measurements were acquired referentially. Custom-made 50 μm (diameter) ultra-low impedance μRC, EIROF electrodes were implanted in dorsal and ventral hippocampus (AP −2.5, ML ±2.0, DV −3.2 mm), (AP −3.9, ML ±2.2, DV −3.1 mm), (AP −5.15, ML −5.35, DV −7 mm), and (AP −6.0, ML 5.0, DV −5.5 mm) to measure hippocampal field potentials. For each hippocampal site, a bundle of two electrodes with ends 125-250 μ staggered apart was used. All coordinates were Bregma referenced.

Hippocampal field potentials were recorded either differentially as local differential measurements, or referentially referenced to a reference screw electrode.

All rats received a lead in the precordium for ECG recordings. The lead is a polyimide flex circuit with a linear array of three 1.5 mm gold electrodes spread 15 mm apart. Electrodes were secured in place with dental cement. The leads were then connected to an electrode interface board (EIB) attached to the acquisition amplifier and all was encapsulated within a 3D printed head-mount. At the completion of the surgery animals were returned to their individual homecages – standard autoclave-ready rat cages with a custom cover that holds the electrical commutator, camera and recording computers – with free access to food and water and maintained at a 12 hour light-dark cycle with lights on between 6 am and 6 pm.

Seven days post-recovery rats were connected to a commutator at the top of the cage via a low-weight USB cable. A single board computer (Raspberry Pi foundation, 3 model B) attached to the cage cover acquired and transmitted data continuously to a network attached storage (NAS). A separate single-board computer (Raspberry Pi foundation, 3 model B) with a low-light level-compatible camera system similarly spooled continuous, time-synchronized video data to the same NAS.

### Data Collection

For mice and rats all biopotentials were acquired at 24 bit resolution and 1 kHz sampling frequency (per channel) via our custom-made data acquisition system that uses the Texas Instruments ADS1299 Biopotential Front End as its core.

The 24-bit digitization provides a dynamic range of 4.5 V with sub-microvolt divisions. Therefore it accommodates the amplitude range and resolution to simultaneously resolve large shifts in tissue potentials (10-30 mV) associated with SD and normal activity in the range of few millivolts associated with local field potentials. This feature complemented with DC-stable μRC electrodes eliminates the need for analog high-pass filtering prior to digitization.

For mice, the acquisition system was designed to provide 8 channels of high quality biopotential recordings. For rats, we extended the acquisition system to provide 16 channels and a 3-axis accelerometer and to fit within a 3D-printed head mounted box (amplifier size: 1“W×1”D×0.25”H).

The raw data contained the information in all frequencies below 500 Hz (the hardware’s digital decimation filter cutoff (Anon, 2017), including bio-potentials from neural activity, spreading depolarization, and electrochemical changes.

### Data Analysis

All recorded data were inspected via in-house written LabVIEW (National Instruments) and MATLAB (MathWorks Inc.) programs that allow for simultaneous re-referencing, filtering, spectral analysis and annotation. For seizure and state of vigilance scoring, the raw ECoG and hippocampal LFPs were band-pass filtered at 1-55 Hz and 1-125 Hz; respectively, to highlight field potentials and seizure dynamics. Electromyogram (EMG) and ECG data were band-pass filtered at 1-125 Hz to extract muscle activity and cardiac dynamics. The three-axis head acceleration data were band-pass filtered at 1-100 Hz.

#### Seizure Detection

For mice, seizures were detected manually from band-pass filtered EEG using custom-written software within LabVIEW environment. Spontaneous seizure activity of more than 10 seconds was identified, scored, and visually verified for origin and evolution pattern according to previously described criteria in (Ssentongo et al., 2017).

For rats, seizures were detected automatically, from band-pass filtered EEG (0.5 < f < 125 Hz) and head acceleration measurements, using in-house custom-written routines in LabVIEW software and verified visually to determine precise onset and end times. Spontaneous seizure activity of more than 10 seconds was marked as a seizure.

#### Spreading Depression Detection

Historically, identifying features of SD have been a large negative shift in tissue potential of order of 10 – 30 mV in amplitude lasting for approximately 30 – 60 seconds (Grafstein, 1956; Kraig and Nicholson, 1978; Somjen et al., 1992). We utilized these features in a semi-automated detection algorithm in a custom written MATLAB (Mathworks Inc.) script to find bouts of SD.

SD events were defined based on negative shifts in tissue potentials greater than 10 mV which lasted longer than 20 s. Onsets and offsets of these events were detected when the derivative of low-pass filtered (< 2 Hz) raw data exceeded four times its standard deviation computed from hour-long blocks of data.

We further visually verified the selected events to achieve temporal accuracy in SD end times which were identified as when DC potential recovered to the pre-seizure baseline. Sequential threshold crossings in spatially adjacent electrodes were then used as indicators of propagation of SD.

#### Included Data

In both animal models of epilepsy, animals were included for further analysis if they had at least one ECoG and one hippocampal depth recordings with sufficient stability. We operationally define “sufficient stability” as having baseline DC fluctuations with amplitudes substantially less than 10 mV over hour-long periods without seizures. All included animals had stable DC recordings throughout their lifetime.

Controls included both animals that received a neurological insult (CM or TeTX) and did not develop epilepsy and ones that did not receive either CM or TeTX.

#### Post-Cerebral Malaria Epilepsy Model

Data were collected from 21 post-CM epileptic mice with 1139 cumulative full days of recordings. Control data included uninfected control animals with no seizures (17 mice with over 820 cumulative recording days), and animals rescued from CM that did not develop seizures (6 mice with 282 cumulative recording days).

#### Rat Tetanus Toxin Model of Temporal Lobe Epilepsy

Data were collected from 6 TeTX epileptic rats with 90 cumulative full days of continuous recordings. Control data included 2 rats with 74 recording days each that did not receive TeTX injections and one rat that received TeTX injection but did not develop epilepsy with 80 days of recording.

#### High fidelity DC-sensitive electroencephalography recording system

In order to capture SDs, sufficient digitization fidelity, including input range and bit resolution plus properly matched electrode impedance to input impedance ratio are required (Hartings et al., 2009; Herreras, 2016; Dreier et al., 2017). Most recording systems have limited bit resolution – typically 16 bits – and therefore impose high-pass filters to limit the input to the range of microvolt to a few millivolt. Our 24-bit digitization – which spans sub-microvolt to volt ranges – does not require such an imposed high-pass filter. Yet especially with small surface area microelectrodes, the electrode polarization could lead to an inherent high-pass filter.

To evaluate the effect of electrode polarization induced by filtering in our recordings we prepared an ionic test system with 2 ionic reservoirs connected by a high-impedance micropipette salt bridge, all filled with a 0.09% NaCl solution (10 times weaker than standard physiological saline). This allows the different chambers to be held at significantly different potentials without driving large charge between them. Each pair of test electrodes was placed into the reservoirs such that individual electrodes were on opposite sides of the salt-bridge. Time-varying potentials (Fig. 1A) between chambers and, therefore between test electrodes, were applied by passing current (Analog Stimulus Isolator, AM Systems Model 2200) between a pair of large platinum plates (0.7×1.7 cm^2^) placed on either end of the salt-bridge (inter-plate distance = 6 cm).

**Figure 1.**
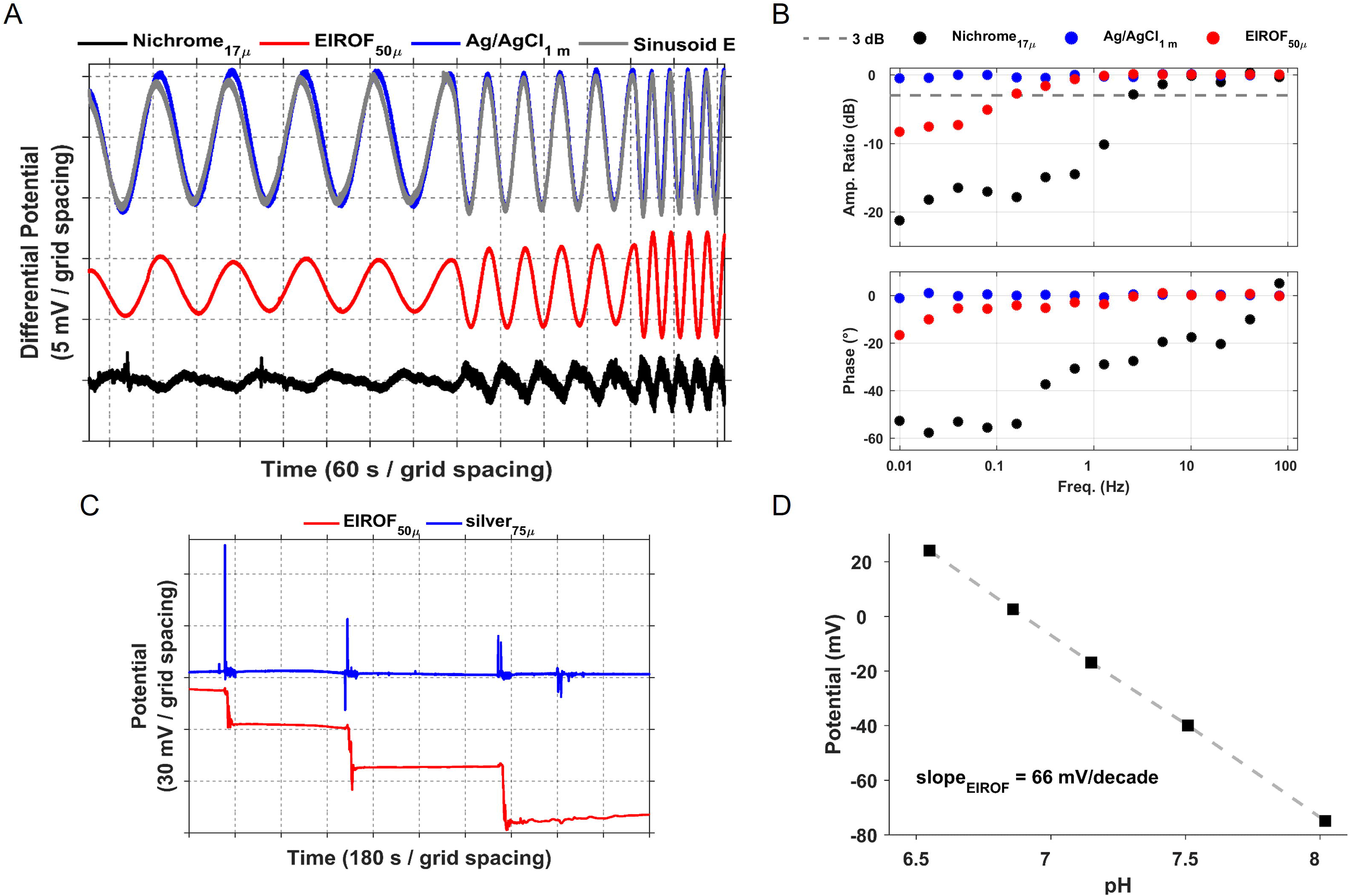
Hardware validation for stable DC and AC measurements. (A) In vitro measurements of phase and amplitude of the electrode potentials across applied near-DC sinusoid electric field. The test electrodes are matched pairs of 50 um (in diameter) microreaction chamber EIROF, 1 mm x 2.5 mm silver/silver chloride pellets (AM Systems, Catalog #550010), and 17 um Nichrome wires placed on one side of a salt-bridge (0.09% saline). Two screw electrodes were used as reference and ground. The sinusoidal electric field with different frequencies was applied via a set of platinum plates on either end of the salt bridge. Test electrodes, reference and ground screws are connected to the acquisition system and from there to a computer through a USB connection. The large Ag/AgCl pellet (blue trace) phase and amplitude follow the sinusoid AC field (gray trace) at all frequencies shown here (0.01, 0.02, 0.04 Hz). While the EIROF electrodes (red trace) show some level of degradation both in phase and amplitude for these very low frequencies, they still sufficiently track the field. However the Nichrome electrodes (black trace) show a large lag and distorted signal. **(B) Low frequency sensitivity of the recording system.** The EIROF can sense and track the field across a wide range of frequencies with only minor attenuation at very low frequencies, in contrast to the much smaller and higher impedance Nichrome electrode that inherently acts as a high-pass filter for the tissue potentials and shows substantial phase and amplitude distortion. **(C)** Unfiltered measurements of EIROF (red trace) and chlorided silver wire (blue trace) electrodes. Each step shift in the potential of the EIROF electrodes indicates an increase in pH. In contrast, Ag/AgCl electrode does not respond to the pH change. The sharp transients in the Ag/AgCl potential timeseries are due to electrode movement and mechanics of titration. **(D)** The potential response of the EIROF electrode to a series of buffer solutions after 1 hour of stabilization.

We found only minor signal attenuation with 50 μm (diameter) μRC EIROF (Fig. 1B red trace) electrodes at frequencies relevant to SD (0.01 Hz). This is in contrast to 17 μm nichrome electrodes (Fig. 1B black trace), which are often used in commercially available tetrodes. The significantly higher impedance of the nichrome electrodes is not well-matched with the input impedance of the amplifier and therefore they inherently create a nonlinear high-pass filter for tissue potentials.

#### pH sensitivity of the EIROF hippocampal depth electrodes

The electrodeposited iridium oxide films (EIROF) are known to be pH sensitive (Papeschi et al., 1976). Physiological brain pH variations occur in the time-scale of seconds to minutes. Therefore, our measurements in the very low frequency ranges are potentially combinations of changes induced by electrochemical reactions (pH) and electrical fields induced by depolarization of neural populations (SD).

We characterized the pH response of the custom-made EIROF deposited 50 μm micro-reaction chamber (μRC) electrodes in the physiologically relevant pH range of 6-8. The electrodes were placed in the Britton-Robinson buffer solution while drop-wise addition of 0.2 M solutions of NaOH and H_2_SO_4_ set the appropriate pH value (measured via a calibrated pH-meter, Anaheim Scientific P771). The test electrodes were all referenced to a pH-insensitive Ag/AgCl pellet electrode (AM Systems, Catalog #550008). The pH-insensitivity of the Ag/AgCl pellet was confirmed via measurements against a double junction Ag/AgCl reference electrode (Beckman Coulter, 3.5 M KCl, Item #A57189). We found the pH sensitivity of our custom-made EIROF deposited μRC electrodes to be 70 mV/log-decade at body temperature of 37°C (Fig. 1C-D).

#### Statistical Analysis

The fraction of seizure-associated spreading depolarization events is the count of seizures with SD from all observed seizures. As estimated rate 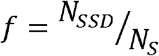, the propagated error Δ*f* for these estimates follow from binomial counting statistics as 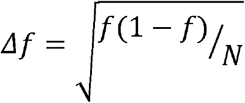 where *N_s_* is the number of all observed seizures, and *N_SSD_* is the number of seizures with SD.

We used non-parametric tests for comparison between different distributions with false positive rate α = 0.01. The distributions pertained to the seizures that ended with SD and those that did not.

When applied to consecutive samples – in particular drawn from different time offsets with respect to seizure termination – each consecutive sample is assumed to be drawn independently from the previous sample. To maintain a group false positive rate for the ensemble of measurements for the given period (0 – 50 s post seizure termination), we rely on binomial statistics. In that, we determine the minimum number of positive tests for the group false-positive rate to also achieve α = 0.01. Reported are the actual count N_true_, maximum count, N_max_, and threshold count N_thresh_. To exclude the possibility that the results are derived from select outlier time-series, we also randomized the labels between the seizure that ended with SD and those that did not. We used 1000 surrogate randomizations and report the maximum count number N_surr_ observed from these.

### Data Availability

All data presented in the figures, including hour-long segments of the EEG, and the code required are available in a shared archive that will become active when manuscript is published.

## Results

### Spreading depression co-occurs with spontaneous, unprovoked seizures

We found frequent instances of spontaneous seizure associated SD in two chronic animal models of epilepsy: the rat tetanus toxin (TeTX) model of temporal lobe epilepsy (TLE) (Jefferys et al., 1995; Sedigh-Sarvestani et al., 2014) and the murine model of post-cerebral malaria (post-CM) epilepsy (Ssentongo et al., 2017).

An example of observed seizure-associated SD in epileptic rat is shown in Fig. 2 A. The seizure onset (sz_start_) and ictal offset (sz_end_) are marked respectively by the red and green dashed line. The seizure onset coincides with an increase in LFP discharges (and is typically preceded by a sentinel spike) and ictal offset is marked as the end of the associated LFP spike-like discharges. The upper panel in Fig. 2A includes high-pass filtered hippocampal LFP and ECoG measurements. We defined the ictal event as the period from the onset of discharges to their offset, including the period of EEG depression in between.

**Figure 2.**
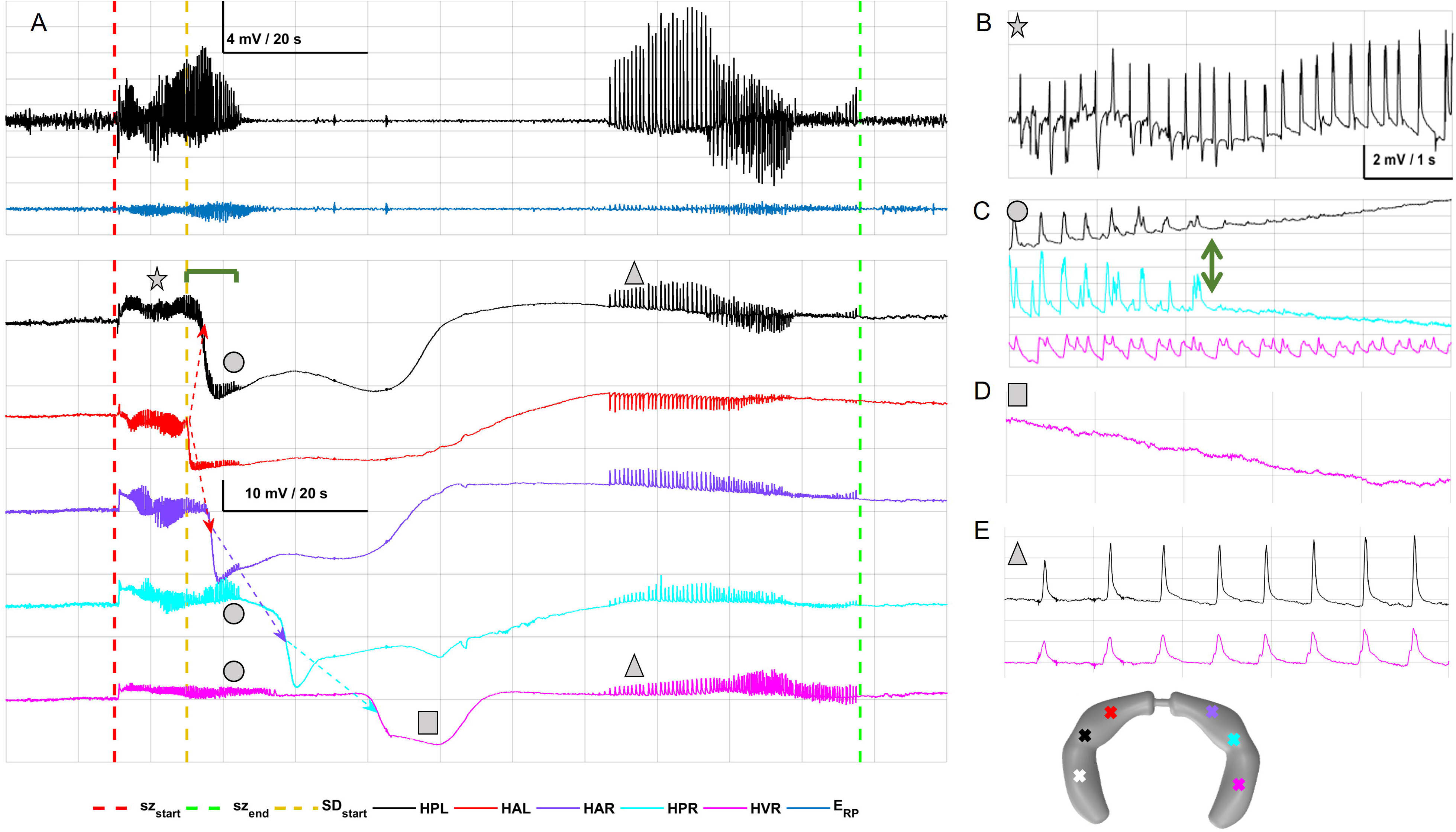
Example of SD from chronic Recordings. Spontaneous seizure with SD-like activity in the hippocampus of a rat under the TeTX model of temporal lobe epilepsy. **(A) Upper panel,** Bandpass filtered EEG, hippocampal (HPL; black trace) and ECoG (ERP; blue trace), highlights the seizure activity with the red dashed line marking its onset (sz_start_) and the green dashed line marking the offset of the ictal event (sz_end_). The EEG depression is evident from the bandpass filtered traces approximately 25 second after the seizure onset. **Lower panel,** Wide-band measurements with the DC component of the signal demonstrate both the seizure activity (B) and the large amplitude slow shift in tissue potential characteristic of the SD (C, D). Although there is minimal residual seizure activity on some channels at the beginning of SD, for the most part the seizure activity terminates during the SD (C, gray star) compared to sites that SD hasn’t reached yet (C, gray square). The EEG suppression is coincident with the negative trough of the SD (D). Dashed arrows indicate the propagation pattern of the SD wave from each hippocampal site. During recovery from SD seizure-like spiking activity appears in all hippocampal sites (E), however during these events the animal shows no sign of behavioral arrest or any other abnormal behavior signatures. The dark green bracket (= 7s) indicates the duration between SD onset (detected from the large DC shift) and EEG suppression in HPL, HAL, HAR, and HPR channels. The onset of EEG suppression in these channels is marked by the double green arrow in (C). Filter settings for traces shown: (A – upper panel) band-pass 1-50 Hz; (A – lower panel, and BE) low-pass at 50 Hz. Channel Names: EPL, ECoG posterior left; EAL, ECoG anterior left; EPR, ECoG posterior right; EAR, ECoG anterior right; HPL, hippocampal posterior left; HAL, hippocampal anterior left; HAR hippocampal anterior right; HPR, hippocampal posterior right; HVR, hippocampal ventral right. The target locations are represented in the schematic of hippocampus in gray. The white cross indicates the area that received the TeTX injection.

The time-matched wide-band recordings sensitive to both seizure and SD are shown in the bottom panel of Fig. 2A. In this example, the SD event, whose onset is marked by the gold dashed line, initiated in a left dorsal hippocampal site and propagated throughout the hippocampi. Similar traces from the murine model of post-CM epilepsy were presented in (Ssentongo et al., 2017 – Fig. 4A). The DC measurements clarify that the period of depressed activity and the later after-discharges are in fact associated with a hippocampal spreading depression event and recovery from it. This phenomenon would not be observable without wideband recordings.

During the pre-ictal period both LFP and DC potentials are at their stable baseline value. As the seizure develops the LFP power increases and remains high during the seizure (Fig. 2B) with complex spike and spike-and-wave signatures in both hippocampal and cortical channels. In
 contrast, DC potentials either remain stable for the entirety of the seizure interval or decrease as the SD initiates mid-seizure (Fig. 2C).

When SD does initiate during a seizure, LFP activity may continue for a short period (4-12 s) following the DC onset on a per-channel basis as illustrated for the HPL channel in Fig. 2C. The high-frequency LFP activity, once SD has initiated, often stops nearly simultaneously (delayed less than a second) across many channels. In the event shown in Fig. 2C, the high-frequency LFP activity ceases across channels in both left and right hippocampus (HPL, HAL, HAR, HPR) approximately 7 seconds *after* the large negative DC shift is observed in the HAL electrode. Utilizing LFP suppression as a marker of SD can therefore lead to missing the correct onset and the spatial origin of the SD event, as well as its propagation.

In seizure-associated SD incidents – regardless of the SD onset time with respect to the seizure – the large negative shift in DC potentials persisted into the post-ictal period coincident with substantial suppression of LFPs (Fig. 2D). At the end of SD, as DC potentials recover to their baseline value, we often observed spike-like discharges in the LFPs (Fig. 2E) after which LFPs also recovered to their initial baseline amplitude. The spike-like discharges shown in Fig. 2E are consistent with the post-SD discharge activity introduced and named as “spreading convulsions” by Van Harrevald and Stamm (Van Harreveld and Stamm, 1953).

Almost all commercial acquisition systems introduce a causal high-pass filter that attenuates and distorts the very low frequencies associated with spreading depression. We illustrate the effect of such high-pass filters on a single hippocampal channel’s signal in Fig. 3. We passed the raw data through 4 pole causal high-pass filters with different cut-off frequencies. As indicated in Fig. 3, evidence of the large negative shift in potential appears for cut-off frequencies less than *f* < 0.5. However, for all added filters the amplitude of this shift is significantly attenuated and there is significant phase distortion in the frequency content so that low-frequencies are also shifted in time with respect to the large amplitude nearly DC shift observed with no high-pass filter.

**Figure 3.**
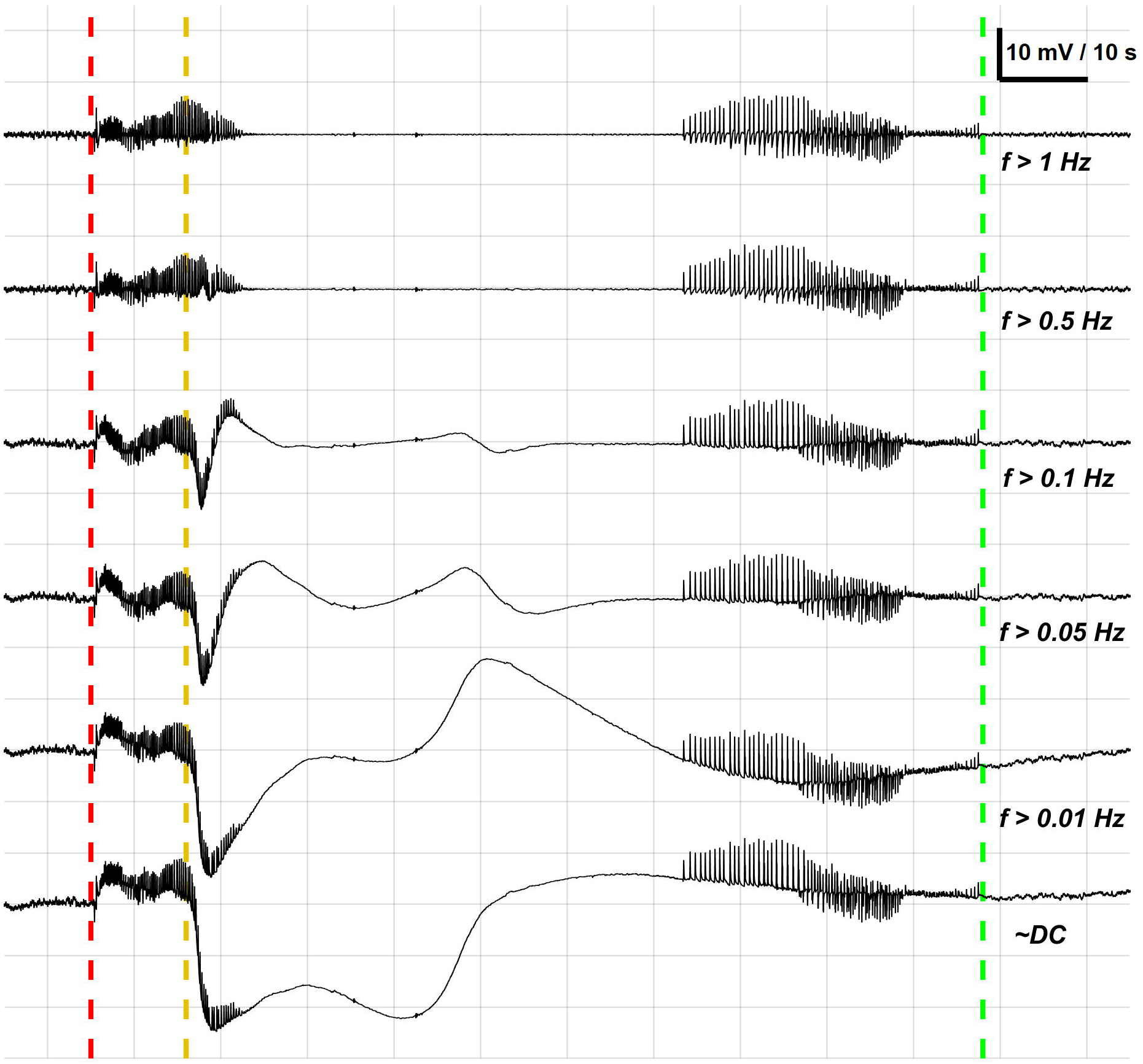
Effect of high-pass filtering on slow shifts associated with spreading depression. Shown is the same HPL channel for the same seizure as Fig. 2 filtered with different high-pass filtering to illustrate the effects of recording system filtering on the character and identifiability of the SD event. Filtered traces were passed through a 4-pole causal high-pass filter with different cutoff frequencies at 1 Hz, 0.5 Hz, 0.1 Hz, 0.05 Hz, and 0.01 Hz. All traces were passed through a causal low-pass filter with cutoff frequency of 50 Hz to conform to finite figure resolution. The long large amplitude DC shift associated with spreading depression is observable in its full detail only when there is no high-pass filter involved. In almost all commercial acquisition systems, a high-pass filter is implemented to accommodate the noise caused by the impedance mismatch between electrodes and the input impedance of the amplifier.

### Large negative shifts in potential are associated with suppression of activity

Characteristic features of SD have been a large shift in tissue potential of order of 10 – 30 mV in amplitude lasting for approximately 30 – 60 seconds (Grafstein, 1956; Kraig and Nicholson, 1978; Somjen et al., 1992). This large and slowly propagating shift is associated with *depression* of activity in extracellular potential. We frequently observed that SD initiated during ongoing seizure activity (e.g. Fig 2A) with seizure-like signatures persisting into the SD event in some channels. Our measurements represent local field potential (LFP) activity that is dominated by synaptic *input* to the local neurons (Herreras and Somjen, 1993a; Herreras, 2016) plus the large scale diffusion potentials that accompany SD (Ryzhkov et al., 2017). Therefore, the seizure-like signatures that continue at the onset of the large DC shifts (Fig. 2C), likely represent nonlocal input from regions still undergoing seizure activity at a distance away from the electrode position.

Classical nomenclature for SD (Leao, 1944) associates a depression of activity that spreads coincident with the shifts in DC potential. To investigate if the DC potentials we observed were associated with suppression of activity, we calculated root mean squared (RMS) of the LFPs over the time interval that included pre-ictal baseline, seizure, and post-ictal/SD activity.

To cover the entire evolution from when the SD event starts while the high-pass filter traces show seizure activity, we selected local field and DC potential values within a window spanning 20 s prior to seizure onset to 20 s after SD offset. The distribution of RMS of band-pass filtered hippocampal LFP (0.5 < f < 125 Hz) in one electrode as a function of the DC potentials of the same electrode is shown in Fig. 4. The data are pooled across all seizures with SD from all epileptic rats. As shown here, when the DC component is large (−12 to −20 mV), the hippocampal LFP activity is typically suppressed. The exception, which broadens these distributions, is that at onset of negative DC shift, seizure activity appears to persist briefly as in Fig 2C. Note that the occurrence of potentials much less than −20mV are much fewer and typically at SD onset, yet the LFP power is still depressed by ~10dB from baseline. This overall correlation of suppressed LFP activity with large negative potential shifts supports that these events are SD.

**Figure 4.**
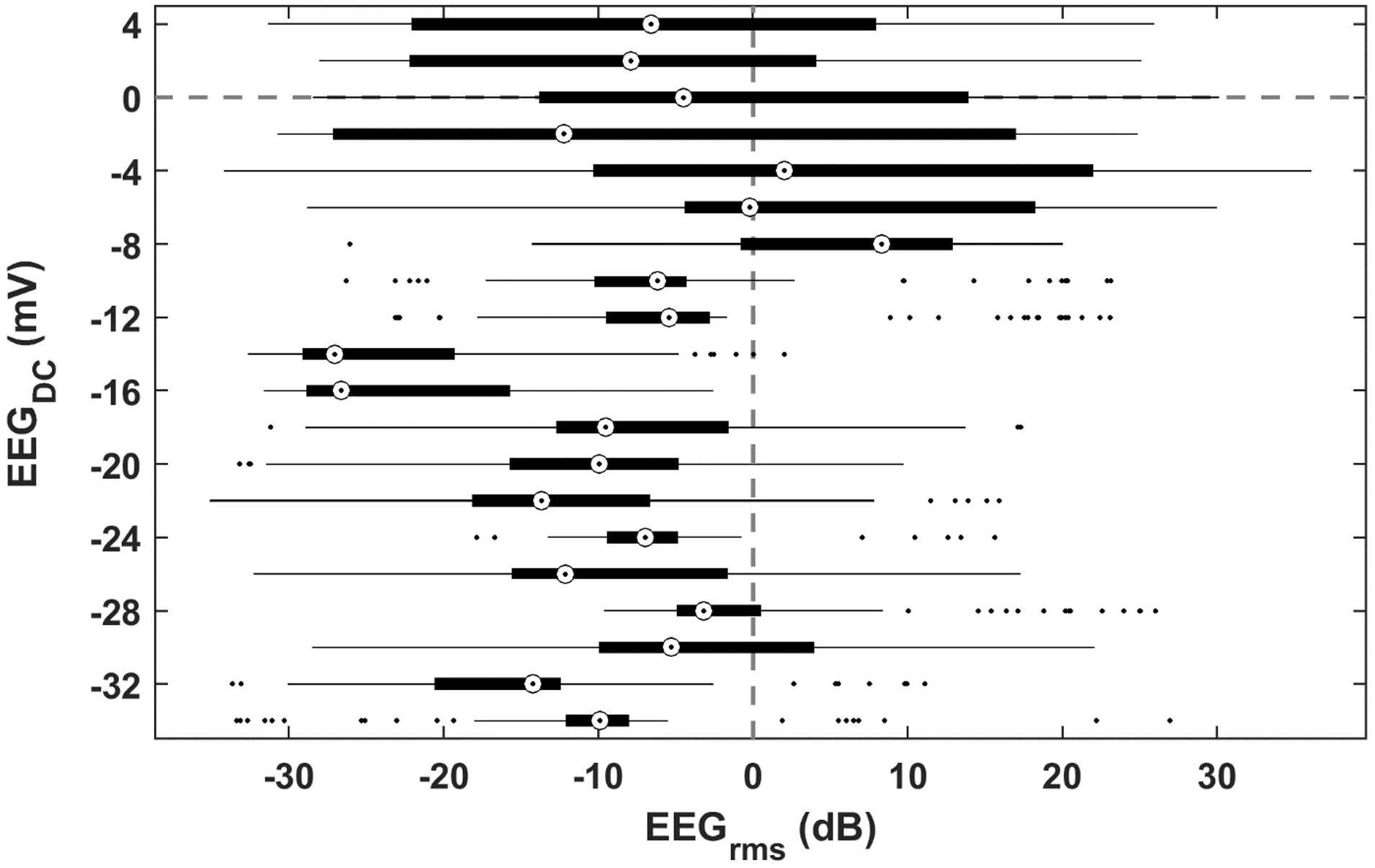
Marked suppression of band-pass filtered EEG during SD trough. Root mean squared (RMS) of the band-pass filtered EEG was calculated over 8 second long overlapping windows in all epileptic animals. Here, we report the RMS values in units of dB; 20*log (RMS/RMS-baseline); where RMS-baseline is calculated as the average of multiple hours of recordings. The RMS of a hippocampal LFP channel was pooled from all seizures with SD. The distribution of RMS values during 20 s prior to seizure onset to 20 s after SD offset where then calculated with respect to the detrended time-matched DC signal. Although as observed in Fig. 2C, sometimes there is residual seizure activity at the onset of SD, the band-pass filtered EEG is fully suppressed during the bulk of the SD. The EEG only recovers to baseline amplitude and frequency after SD recovery.

### The signature of spontaneous seizure-associated SD in two animal models of epilepsy

The two animal models of epilepsy utilized in this study are fundamentally different. The TeTX model of temporal lobe epilepsy is induced with intrahippocampal injection of tetanus toxin. This creates a focal lesion with modulated neuronal excitability, and produces secondary generalized seizures. In this model seizures emanate from sites in the ventral hippocampi and generalize to other hippocampal and cortical regions.

The murine model of post-CM epilepsy mimics human conditions of post-infection acquired epilepsy with a long period of epileptogenesis (Bahari et al., 2018) followed by spontaneous unprovoked seizures. In this model the brain undergoes widespread multifocal damage, and the seizure origins and evolution patterns vary.

Despite the pathophysiological differences, in both models we observed hippocampal SD with occasional instances of SD invading cortex (data not shown). SDs often initiated during ongoing seizure activity (Fig. 2A n = 410 from 425 SDs in rats and n = 99 from 172 SDs in mice). We also observed instances where SDs occurred shortly after recovery from seizures (n = 15 from 425 SDs in rats) and (n = 70 from 172 SDs in mice). In epileptic rats, we found 21 seizure clusters comprising a total of 56 seizures. These events were associated with 124 of 425 total SDs.

The seizure-associated SD shown in Fig. 2A is representative of multiple incidents (n = 143 from 301 seizure associated SDs outside of seizure clusters in rats and n = 3 from 172 SDs in mice) where SD recovery to baseline value was accompanied by seizure-like spiking activity. This pattern of spikes, shown in Fig. 2E, is consistent with recovery dynamics from acutely-induced SD that are termed spreading convulsions (Van Harreveld and Stamm, 1953; Lauritzen et al., 2011).

Illustrated in Fig. 5 are details of recording duration, seizure and SD occurrence, seizure origin, and type of death for each epileptic rat (Fig. 5A) and mouse (Fig. 5B). In the murine model of post-CM epilepsy SD was observed in all four mouse-parasite strain combination studied. In this model seizure initiation and evolution varies within and across animals. Within the coverage of our electrodes however, we found seizure-associated SD predominantly in hippocampus. We extended the electrode coverage in the TeTX rat model of TLE.

**Figure 5.**
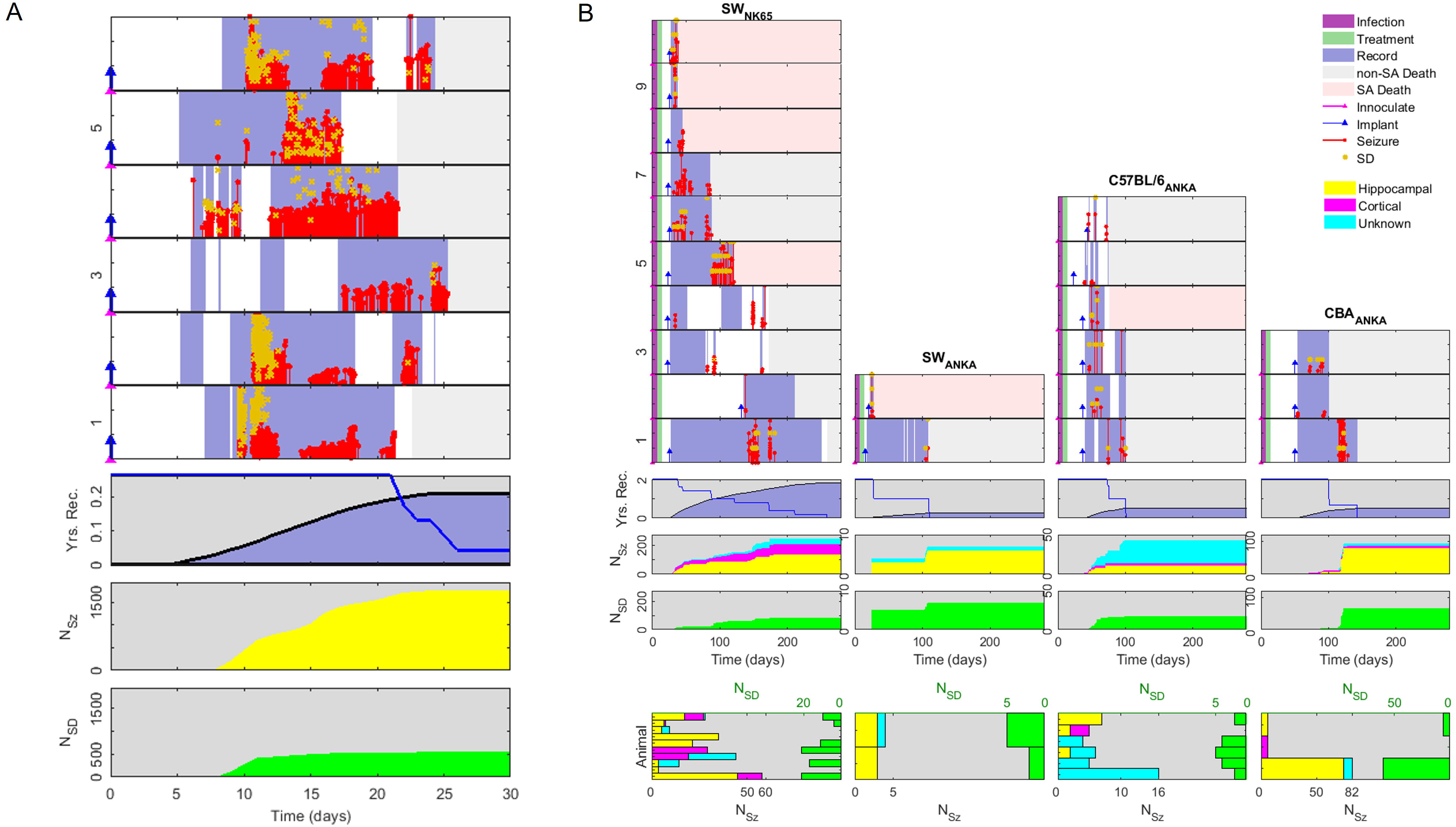
Time course of the experiment from inoculation through to death for rats (A) and mice (B). In the upper panels, time courses of all recorded animals, with columns for experimental strain combinations (for mice), and row for individual animals. The time courses begin with inoculation (magenta triangle) at day 0. The time of electrode implantation is indicated with blue markers and the period of recording is marked in blue background. For mice, the infection and antimalarial treatment periods are marked by the purple and green backgrounds. The time of seizure occurrence over (seizure length > 10 s) is indicated by red markers, with height representing duration in minutes. Time and duration of SD events are indicated with gold markers. Time of death is indicated by the transition to grey (from investigator sacrifice, nonseizure associated death) or pink (seizure associated death) shading. The lower panels represent cumulative time of recordings in blue and uncensored survival curves (blue trace), cumulative number of seizures observed, and cumulative number of SD events observed. Total recording time represented is 90 for rats and 1139 days for mice. The cumulative number of seizures (>10□s long) marked is 441 for mice and 1256 for rats, with none observed from control animals. For rats the seizures all have hippocampal origin. For mice seizures were classified by their origins into hippocampal (yellow), cortical (magenta), and unknown (cyan) (Ssentongo et al. 2017).

Collectively SDs were associated with approximately one third of the seizures in each animal model. For seizure clusters, we only counted the first seizure as the representative.

In the TeTX rat model of TLE, we observed seizure-associated SD in all 6 of 6 animals we report here. We observed an overall seizure-associated SD rate of 33% ± 1% during 90 cumulative days of recording from 1256 total seizures. Note these counts include seizure clusters with multiple SD events. If we only count the first seizure and its associated SD event for each seizure cluster, the overall seizure-associated SD rate would be 26% ± 1% (322 events out of 1221 total seizures).

The TeTX rat model of TLE has relatively small variability in latency to first seizure (6-12 days), and a seizure rate that peaks early to many seizures an hour in the following week, and then wanes to one seizure every few days after a few weeks. In this model we observed more frequent SDs per seizure during the higher seizure rates (Fig. 6A and C). Note that we started recordings late due to technical difficulties for the one rat in which we observed the fewest SD events. That rat (number 3 in Fig. 5A) presented 3 SD events during the recording period.

In this model, seizures typically emerge from the hippocampus close to the toxin injection cite, and seizure-associated SD events most frequently emerged from near the same focal point and spread.

In the post-CM epilepsy model, we observed multiple instances of seizure-associated SD in 15 out of all epileptic mice under study (6/10 SW-NK65, 2/2 SW-ANKA, 5/6 C57-ANKA, 2/3 CBA-ANKA). Across 1139 cumulative days of recordings from 21 epileptic mice the estimated seizure-associated SD rate was 38% ± 2%.

The seizure origin and evolution patterns in the post-CM epilepsy model was more complicated with a very wide distribution of latency to first seizure, and periods of seizures that appeared interspersed with long (days to weeks) seizure free periods (Fig. 5B). The large variation in latency makes interpretation of variations of the slope of the cumulative SD count (Fig. 6B) more difficult to interpret.

**Figure 6.**
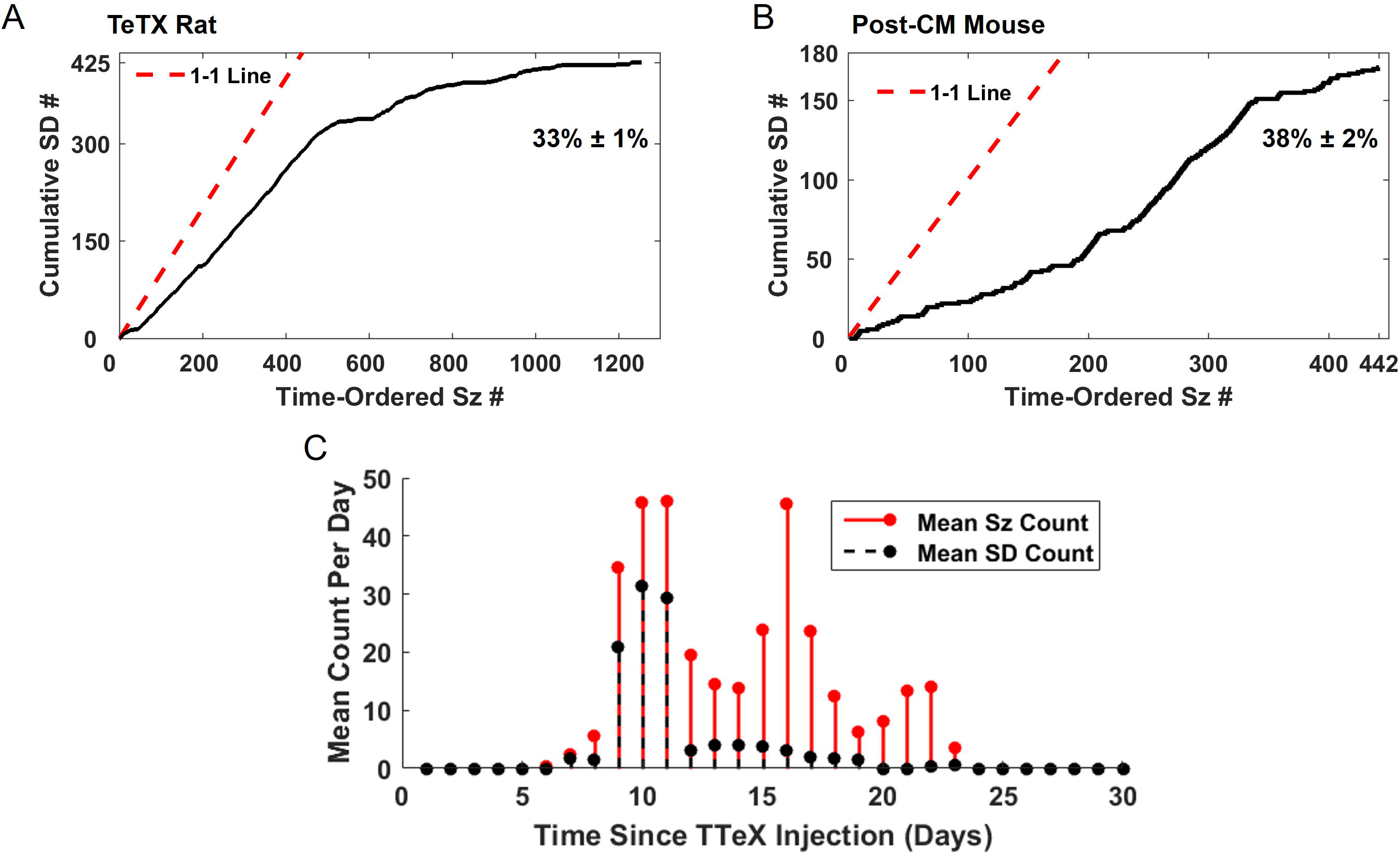
SD is associated with a substantial percentage of seizures in both animal models. **(A, B)** Cumulative count of seizure-associated SDs as a function of seizures rank-ordered by time of occurrence in (A) TeTX rat model of TLE and (B) post cerebral malaria mice. Overall, 33% of the 1256 collected seizures in epileptic rats had associated SDs, and 38% of the 441 collected seizures in epileptic mice were associated with SDs. Estimated propagated error rates are 2% or less (see methods). **(C)** Mean seizure and SD rate per day per animal averaged over all epileptic animals in the rat TeTX model of TLE for the days they were recorded. The model has a high seizure rate during weeks 2 and 3 post-induction. During this time the SD per seizure rate is also high.

### Propagation Velocity of observed SD events

In order to estimate the propagation speed of SD in chronically epileptic tissue, we calculated the Euclidean distance between electrodes implanted in hippocampi. In post-CM epileptic mice, SD propagated between left and right hippocampus with an estimated propagation speed of 6 ± 1.2 mm/min (mean ± std). This is consistent with in vitro (Aitken et al., 1998) and acute induced in vivo SD experiments (Herreras and Somjen, 1993b, 1993a; Somjen, 2004; Aiba and Noebels, 2015; Loonen et al., 2019).

In TeTX rats, apparent propagation speed was calculated and averaged over an ensemble of seizure events that displayed a stereotyped behavior of SD during non-clustered seizures that initiated at the position marked HPL in Fig. 7A and spread through both hippocampi (n = 299 from 425 total SDs). In this case, SD appeared to be transmitted between adjacent hippocampal electrodes with mean apparent speeds ranging from 6 to 20 mm/min within left or right hippocampi (Fig. 7A), and much faster apparent propagation between hippocampi (Fig. 7B).

**Figure 7.**
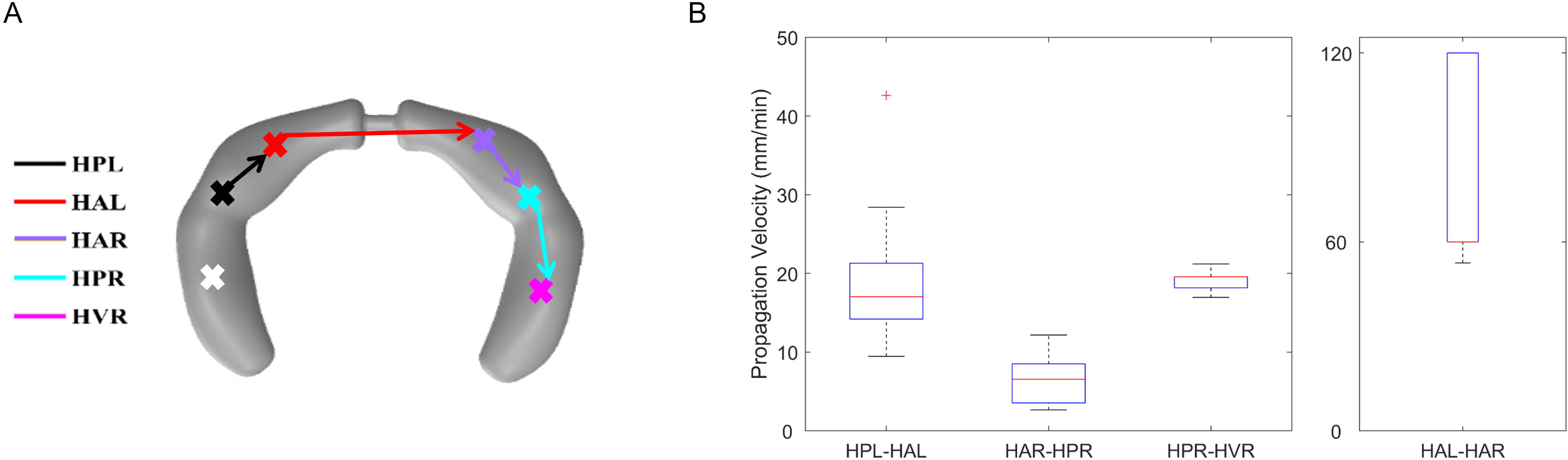
Apparent propagation speed of hippocampal SD in TeTX rat model of TLE. We calculated the time in between appearance of consecutive SDs in electrodes and divided it by the Euclidean distance between the electrode sites. **(A)** Illustration of electrode positions and path steps: left posterior hippocampus (HPL) to left anterior hippocampus (HAL); black arrow. HAL to right anterior hippocampus (HAR); red arrow. HAR to right posterior hippocampus (HPR); purple arrow. HPR to right ventral hippocampus (HVR); cyan arrow. **(B)** Distribution of speed for each path step. The speed distributions are mostly within ranges previously presented in the literature (<20 mm/min). However, the apparent speed is much faster for the left to right propagation (HAR-HAL). The white cross in (A) indicates the TeTX injection site.

### SD did not occur in control and non-epileptic animals

In order to confirm that SD was associated only with seizures, we scored data from age-matched control animals with sufficiently stable measurements. In rats controls included animals that did not receive TeTX injections (N = 2, 154 cumulative recording days), and animals that received TeTX injection but did not develop spontaneous seizures (N = 1, 80 recording days). In mice, control data included uninfected animals with no seizures (N = 17, 820 cumulative recording days), and animals rescued from CM that did not develop seizures (N = 6 mice, 282 cumulative recording days). We found no SD events in any of these control groups, which indicates that the SD was physiological – not an artifact of the recording system – and caused by the epilepsy.

### Hippocampal SD is associated with suppressed cortical activity

In order to investigate if hippocampal SD modulates cortical activity, we calculated normalized power of one hippocampal LFP lead (HVR) and one ipsilateral cortical lead (right posterior ECoG) for a time window spanning from pre-ictal to post-SD recovery times. The data from all TeTX rats across all non-clustered seizures were divided into two groups: seizures that ended in SD and seizures without SD. Normalized spectral power (within frequency band of 0.5 – 75 Hz and 0.01 normalization factor) was computed as a function of time with respect to seizure offset, or SD onset, for each seizure.

The time dependency of the average power across seizures within each group is shown in Fig. 8A. Both the hippocampal LFP and ECoG power decrease post-ictally from their ictal values. As expected from Fig. 4, during SD the average post-ictal power in the hippocampal LFP is lower than the comparable average power post-ictally without SD (Fig. 8A, upper panel). A similar decrease in cortical spectral power is observed during hippocampal SD when compared to post-ictal periods without SD (Fig. 8A, lower panel). Note that SD, as represented by large DC potential changes, did not propagate to the cortex during any of these rat recordings.

**Figure 8.**
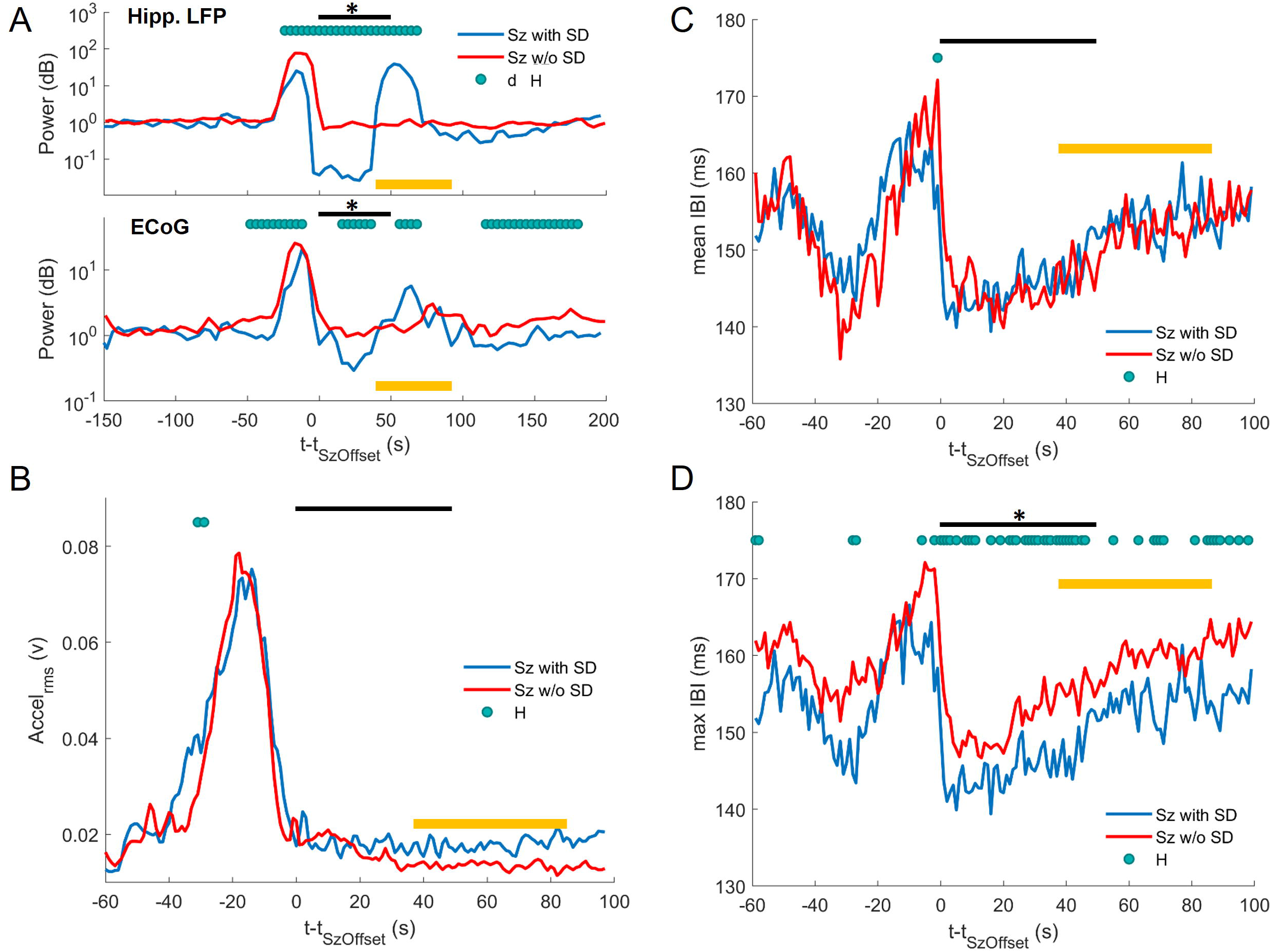
Peri-SD Onset vs Peri-Seizure Offset Metrics in epileptic rats: (A) EEG Power in hippocampal LFP and ECoG, (B) Movement, and (C, D) Cardiac Dynamics. Shown in each case are the means of the time-resolved distributions of each metric (solid lines) and indicator of significantly different distributions (green marks, Wilcoxon ranksum test, alpha = 0.01), where events were aligned at the end of seizures and separated into those that ended with SD (blue lines) and those that ended without SD (red lines). **(A)** Hippocampal LFP and ECoG power with respect to seizure-offset time averaged over all seizures with SD and without SD. In ictal events with SD, the hippocampal LFP demonstrates a significant decrease in power post-seizure offset compared to those without SD. This is coincident with SD events. Similar, although smaller, decreases in the ECoG power were observed. Power is calculated over 4-second long overlapping windows with 2-second overlap. Within the first 50 seconds (indicated by the black line), given the overlapping windows and binomial statistics, we need at least 2 repeated observations of difference between power in the two groups (with and without power) for the difference to count as statistically significant (marked by the black star). We observe 13 for hippocampal LFP and 6 for ECoG. **(B)** Head acceleration with respect to seizure-offset time averaged over all seizures with SD and without SD. The seizures in the TeTX rat model of TLE are convulsive. At the beginning of the seizure animals show head-bobbing and freezing behavior and full-body convulsions start towards the second half of the seizure. This is shown via the head acceleration as it peaks approximately 20 seconds prior the seizure offset for both groups (seizures with SD and seizures without SD). The Wilcoxon ranksum test (alpha = 0.01) was performed for individual time-points and showed only two points are significantly different from one another, marked by the green dots (H). Similar binomial statistical analysis as that of the EEG confirmed that the groups are not statistically different from one another. The yellow line indicates the time where in the EEG we observe spreading convulsion. **(C)** Mean inter-beatinterval (IBI) with respect to seizure-offset time averaged over all seizures with SD and without SD. The mean IBI is calculated in overlapping windows (window length = 2s, overlap = 1s). Both groups indicate longer IBIs towards the end of the seizure. The Wilcoxon ranksum test (alpha = 0.01) confirmed that the groups are not statistically different from one another except in one point, marked by the green dots (H). **(D)** Max inter-beat-interval (IBI) with respect to seizure-offset time averaged over all seizures with SD and without SD. The max IBI is selected in the calculation process for the mean IBI in (B). Although the mean IBI does not show any significant difference between the two groups, the max IBI for seizures with SD is significantly shorter compared to seizures without SD (Wilcoxon ranksum test, alpha = 0.01) for most of the time (green dots). Calculations based on binomial statistics confirm that the number of times (n=30) these time points are different from one another is outside random occurrences.

To quantify if the spectral power was statistically different between the two groups, we compared the distributions of spectral power within groups as a function of time using Wilcoxon Rank-Sum test. Marked in Fig. 8A with green dots are all time points that the groups are significantly different from each other (α = 0.01). These positive test patterns mimic the difference indicated by the differences in means. The number of positive detections are far more than would be expected from binomial statistics with false positive rate *α* = 0.01. To test if the positive detections could be attributed to a small fraction of the SD events, we compared results with 1000 surrogate random permutations of the labels. The number of time points observed with correct labels far exceeded that of the surrogates.

### Hippocampal SD is not associated with overt behavioral changes

SD is known to affect cognition and consciousness (Eikermann-Haerter et al., 2013). This has led to hypotheses on the role of SD in postictal amnesia and loss of consciousness (Gorji, 2001; Butler and Zeman, 2016). In order to determine whether SD has an effect on overt animal behavior, we looked to see if seizures with SD caused loss of consciousness or were accompanied with any obvious lack of motion. We measured root mean square (RMS) of head acceleration (Accel_rms_) as a function of seizure offset time. Shown in Fig. 8B is the average of Accel_rms_ over all seizures without SD vs the average of Accel_rms_ over all seizures with SD from 60 s prior to seizure offset to 100 s after. Comparisons of the time-dependent distributions of Accel_rms_ between groups (Wilcoxon Rank – Sum, *α* = 0.01) are also marked with green dots if significantly different.

We found no significant change in head acceleration due to existence of SD. From time-matched video recordings we confirmed that SD often occurs when the animal is already recovered from the seizure and is moving normally. Further, the discharges during spreading convulsion (indicated by the yellow line in Fig. 8) were not accompanied by any sign of abnormal activity in the accelerometer measurements. This was consistent with visual inspection of the time-matched video. From these measures, the animal’s overt behavior in the post-ictal period appeared independent of the presence of SD and distinct from standard ictal behavior.

### Hippocampal SD is associated with autonomic nervous system changes

To address the effect of SD on autonomic regulation, we quantified cardiac activity as a function of time with respect to seizure offset or SD onset. ECG was first analyzed to identify each heart beat and its associated inter-beat interval (IBI). Within 2 s long windows the mean and maximum inter – beat interval (mean IBI, max IBI) were computed. Averages of the mean IBI for seizures that end without (red) and with (blue) SD as a function of time from seizure offset are presented in in Fig. 8C. Non-parametric test of differences in distributions of mean IBI (Wilcoxon Rank-Sum, *α* = 0.01) confirms that these distributions are not substantially different, with few green indicators shown.

The averages, and distributional comparison indicators, for the max IBI across seizures that end without (red) and with (blue) SD are shown in Fig. 8D. Although the distributions of mean IBIs are not different between these groups, the distributions of max IBIs are substantially different.

The statistics here are similar to that performed for power spectrums in Fig. 8A. The positive detections are far more than what is expected from binomial statistics with *α* = 0.01 and from surrogates analysis in which the labels are randomly shuffled.

### SD may modulate brain susceptibility to future seizures

In the rat TeTX model of TLE we observed multiple seizure clusters (n = 21 with total of 56 seizures) where 2 or more seizures occurred within a 10 minute interval. Each of these cluster events was initiated with a seizure followed by a SD event. During clusters, SDs appeared to connect what otherwise appear as individual seizures.

What would appear from the AC recordings as separate seizures with relatively short interseizure interval is shown in the upper panel of Fig. 9A. The connection between these events is only evident from the DC component of the recordings highlighting extensive SDs (Fig. 9A bottom panel). The first seizure shown is preceded with SD initiated in the HPR channel, and additional SD events bracket and interplay with the subsequent seizure 400 s later. The event shown in Fig. 9A comes from the middle of a seizure cluster that started with a seizure 413 s earlier. Only in seizure clusters did we find SD occurring prior to a seizure, and did so only for secondary seizures within the cluster. In other independent single seizures with SD (n = 301), in the epileptic rats, SD occurred either during or after the seizure.

**Figure 9.**
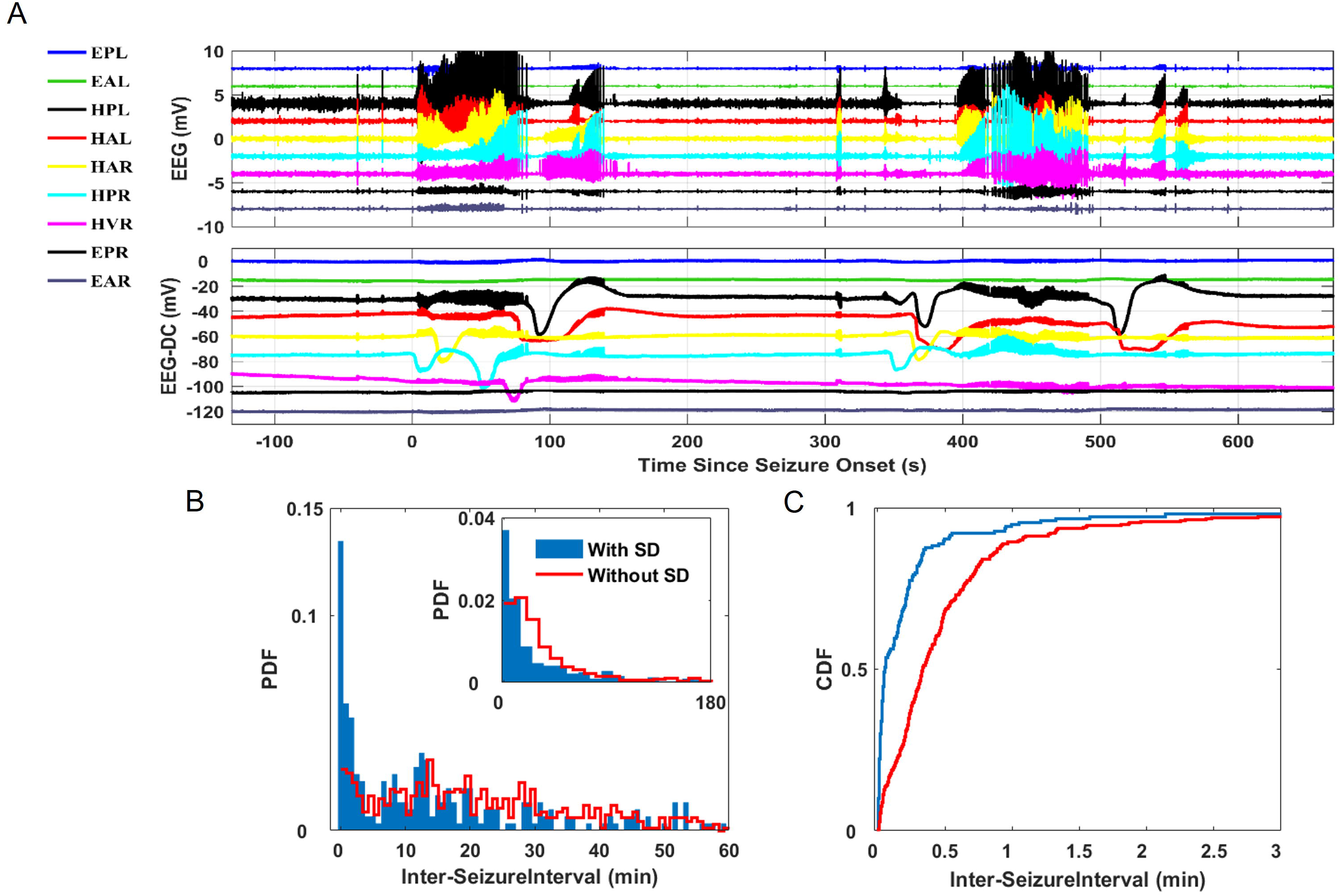
SD appears to mediate seizures in seizure clusters in the TeTX rat model of TLE. **(A)** Example of a pair of seizures within a seizure-cluster. Band-pass filtered LFPs (upper panel, filtered 1 < f < 50Hz) and full-band measures are shown for two seizures that occur within 5 minutes with SDs occurring during and in between the seizures. Note that this cluster had started earlier with a previous seizure ending 320 seconds before, and that the first seizure shown here starts coincident with an SD event. **(B, C)** Inter-seizure interval (ISI) probability density (B) and cumulative distribution (C) for inter-seizure intervals with and without SD. There is a dominant peak at very short intervals for ISIs with SD that does not appear for ISIs without SD.

To characterize the effect of SD on seizure interplay, we quantified the inter-seizure interval for seizures with and without SD (Fig. 9B-C). We found that inter-seizure interval for seizures with SD was significantly (Kolmogorov-Smirnov test, p = 1.25e-6) shorter than for seizures without SD. The distributions were notably different for inter-seizure intervals less than 5 minutes (Fig. 9B), therefore associated with seizure clusters, with a large peak in inter-seizure intervals below 5 minutes for intervals that include SD.

### Observation of Cyclic SD

Recurrent or cyclic SD was first demonstrated in focal ischemic brain as a result of occlusion of middle cerebral artery in rats (Nedergaard and Astrup, 1986). Since then it has been reported in many clinical recordings from patients with aneurysmal subarachnoid hemorrhage (Dreier et al., 2006; Oliveira-Ferreira et al., 2010; Chen and Ayata, 2016), intracranial hemorrhage (Lauritzen et al., 2011; Hertle et al., 2012), traumatic brain injury (Fabricius et al., 2006; Hartings et al., 2009, 2014), and migraine (Dreier et al., 2005).

We observed an incident of cyclic/recurrent SD following a generalized seizure in one epileptic rat (Fig. 10A). In this event we observed seizure-associated SD in the left and right ventral hippocampus followed by another episode of SD in the right ventral hippocampus approximately 200 s later that was not otherwise associated with additional seizure activity.

**Figure 10.**
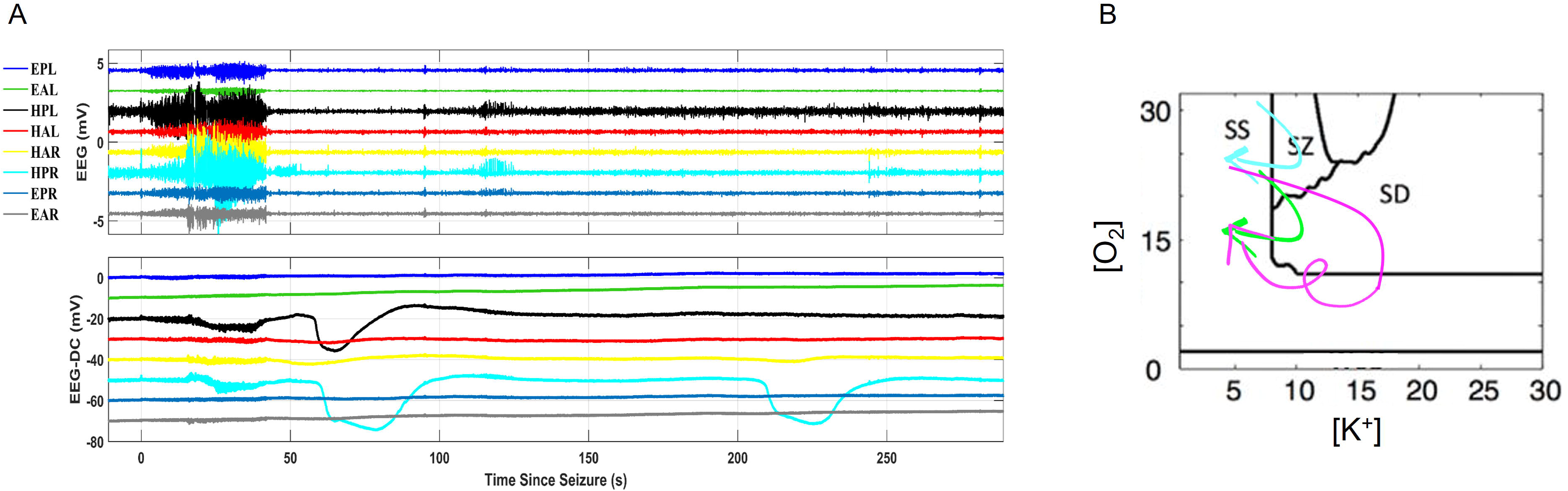
Recurrent SD in time and state space: **(A)** Example of a seizure that is punctuated with a SD event that is recurrent in one channel without additional seizure dynamics. Shown are band-pass filtered (top traces, 1 < f < 50Hz) and full band (lower traces) field potentials. Channel names: EPL; ECoG posterior left, EAL; anterior left, HPL; hippocampal posterior left LFP, HAL; hippocampal anterior left LFP, HAR; hippocampal anterior right LFP, HPR; hippocampal posterior right LFP, HVR; hippocampal ventral right LFP (contralateral side to the tetanus toxin lesion), EPR; ECoG posterior right, EAR; ECoG anterior right. **(B)** Two-parameter state space based on extracellular oxygen and potassium concentrations adapted from (Wei et al. 2014) separated into computationally prescribed regions of steady state (SS), spreading depolarization (SD), seizures (SZ), and tonic firing (TF). The cyclic SD event shown in (A) could correspond to a trajectory in this state space as illustrated by the magenta arrow. Other observed types of seizure/spreading depression dynamics observed in the rat model are also illustrated: seizures with no SD correspond to cyan trajectory; and seizures followed by SD would correspond to the green trajectory.

## Discussion

We recorded spontaneous seizure-associated SD in two different animal models of chronic epilepsy using high dynamic range and low-frequency stable recordings collected over weeks to months per animal. The stability of the recordings permitted us to monitor SD and its propagation with respect to spontaneous recurrent seizures in freely behaving animals. In these models, SD is associated with more than a third of all seizures, and appears to connect seizures in seizure clusters.

The two animal models in this report are both models of acquired epilepsy but with very different origin and mechanism. The TeTX rat model is induced by a focal injection of tetanus toxin that is then retrograde transported and passed across synapses where it cleaves and interferes with SNAP complexes of inhibitory neurons, leading to an approximately focal hippocampal seizure onset zone without significant early neuronal dropout or morphological changes in the tissue. Latency to first convulsive seizure is fairly restricted to 8-12 days post injection, and the majority of seizures emerge from near the focal injection point. The postcerebral malaria mouse model is induced by a bout of cerebral malaria, which acutely is marked by widespread vascular blockage and extension, micro-hemorrhages and brain edema. Long term this results in significant morphological changes, including neuronal cell dropout, expanded ventricles, and hippocampal and cortical sclerosis (Ssentongo et al., 2017). Latency to first convulsive seizure is varied ranging within 27-120 days, and seizures appear to originate from hippocampus, cortex, or with undetermined origin even in the same animal (Ssentongo et al., 2017). In this model the SD events were uncorrelated with seizure origin, and were primarily observed in hippocampal leads. Because of the significantly different pathophysiology of these models we conclude that the emergence of SD from seizures is not likely to be linked to the specific mechanisms underlying the seizure susceptibility of these epilepsies, and assert that SD is likely a frequent phenomenon in the chronically epileptic brain.

Spreading depolarization has been reported recently both in clinical settings from human EEG acquired from patients with low mobility in intensive care units (Fabricius et al., 2006, 2008; Drenckhahn et al., 2012) as well as in awake animals (Khoshkhoo et al., 2017). In these cases SD is defined as a propagating event with rapidly developing reduction of EEG power which is then followed by gradual recovery (Strong et al., 2002; Fabricius et al., 2006, 2008; Khoshkhoo et al., 2017). The SD events in these studies are determined either by negative shifts in near-DC ECoG measurements with amplitudes of 1-3 mV and durations of 1.5-3 minutes (Fabricius et al., 2006, 2008), or by EEG power suppression (Khoshkhoo et al., 2017).

Our high-pass filtered time-series of the SD event in freely behaving epileptic animals (Fig. 2A and Fig. 3) mimic these characteristics. However, as shown in Fig. 2A and Fig. 3 using high-pass filtered data or EEG power will lead to a significantly delayed detection of the SD onset, offset, and propagation pattern. In the example shown in Fig. 2, the EEG suppression starts 7 s *after* we detected SD onset from the DC measurements (dark green bracket in Fig. 2A). Because the EEG suppression appears almost simultaneously across 4 of the hippocampal channels (HPL, HAL, HAR, HPR), it is not reflective of the propagating depolarization event. Our findings therefore underscore the value of DC-sensitive stable measurements to investigation, diagnosis, and control of SD.

We defined the ictal event as the period to include the first set of discharges (classic electrographic seizure), the period with large low-frequency potential shift and associated EEG depression (SD), and the recovery after-discharges, i.e. spreading convulsions (Fig. 2). In the paper “Spreading cortical convulsions and depressions” Van Harrevald & Stamm reported spike activity following cortical SD and were first to coin the name “spreading convulsion” for the discharge, or “convulsive”, activity they observed in the cortical measurements (i.e. Fig. 3 in (Van Harreveld and Stamm, 1953)). In their seminal work the term “spreading convulsion” was used to refer to a phenomenon in the EEG with no mention of motion measurements i.e. muscle jerks. Likewise, we did not observe any behavioral or motion abnormalities during recovery from SD or spreading convulsions (as shown in Fig. 8B).

As shown in Fig. 2A and Fig.3, in the presence of only high-pass filtered data, seizures followed by SD and spreading convulsions appear as two *separate* seizure periods with a period of EEG suppression between them. The wide-band recordings uncover the SD event and confirm that these events are indeed not separate but are part of one ictal event. Suppression of EEG amplitude during seizures have been reported both in animal and human studies (Cook et al., 2016; Smith et al., 2018). Smith et al. reported ictal events comprising spiking activity periods often interdigitated with suppressed EEG intervals which they defined as mid-ictal and post-ictal depression. These events can likely be explained by an underlying SD. But without DC-sensitive recordings, depressions in activity such as the period 30 s into the seizure in Fig. 9A (HAR channel) could not be definitely attributed to SD. Therefore, it is likely that our observations extend beyond the two models we describe here.

The realization that ictal events might also include SD phenomena opens new targets for epilepsy treatment development. On the other hand, understanding that the occurrence of SD might lead to inaccurate seizure counts allows for better screening of anti-epileptic drugs for seizure control. In addition to treating seizures, control of SD occurrence may lead to better outcomes, fewer seizure clusters, and what is now considered postictal phenomena such as memory deficits and headaches.

We used a commercially available and human compatible EEG-analog front end (ADS1299, Texas Instruments) with sufficiently non-polarizing microelectrodes to achieve high fidelity at frequencies below 0.01 Hz. Many clinical EEG acquisition systems routinely used in epilepsy monitoring units utilize similar core amplifier hardware and are matched to low-impedance electrodes. Therefore, translation of our findings into clinical settings only requires firmware and software adaptations. Such recording platform would allow the community to investigate spontaneous SD in epilepsy and adapt diagnosis and treatment appropriately.

Clinical findings from patients with subarachnoid hemorrhage (Dreier et al., 2012) or acute brain injury (Fabricius et al., 2008) identify observed SDs as a risk factor for later development of seizures. But these reports significantly correlate SD frequency and duration with increased acute measures of damage such as infarct size and related symptoms such as fever and low blood pressure. These results therefore imply that subsequent epileptogenesis is more likely a consequence of tissue damage and that acute SD is a secondary measure of that acute damage. Our observed link between high seizure rate and high SD rate might imply that the same phenomena that make the tissue susceptible to seizures also makes it susceptible to SD. But the relationship in absence of spontaneous seizures is complex. Reports have suggested both higher and lower tissue resistance to induced SD in *in vitro* studies of animal epilepsy models and human epileptic tissue (Köhling et al., 2003; Gorji and Speckmann, 2004; Petzold et al., 2005; Maslarova et al., 2011) as well as in awake rats with pentylenetetrazol (PTZ) induced seizures (Koroleva et al., 1993). Our findings that SD only occurred during seizures or linking seizures in seizure clusters is more consistent with the idea that the seizure itself transiently makes the tissue susceptible to or induces SD.

The mechanistic relationship between *spontaneous* seizures and SD remain to be experimentally investigated. Computational modeling of a single-compartment Hodgkin-Huxley style model of single neuron and extracellular space suggest a unification between steady-state, spikes, seizures, spreading depolarization, and the depolarizing wave of death (Wei et al., 2014a). In an updated version of the model cellular volume fraction is shown to orchestrate brain state trajectories from seizure through spreading depolarization dynamics (Ullah et al., 2015).

Our findings support the predicted trajectories of the Ullah et al. model. Our measurements can be mapped to model states of steady state (SS), seizure (SZ), and SD. Our observed transitions (as shown in Fig 2, 8A, and 10B) between these states are represented by continuous trajectories through the model’s state space illustrated in Fig. 10A. Even the rare *in vivo* observation of cyclic/recurrent SD as presented in Fig. 10B is readily described by the predicted trajectories of the model (looping magenta line). Therefore, we propose that the model contains sufficient relevant physiological dynamics to inspire new targeted experiments to investigate the underlying mechanisms of seizures and SD as well as to control their evolutions.

The variables represented in the Ullah et al. model that modulate SD and seizure susceptibility, including extracellular volume and tissue oxygenation, will also be modulated *by* SD and/or seizure incidence. These variables are shown to have much longer recovery dynamics than the SD and seizure events (Olsson et al., 2006; Farrell et al., 2016).

In our observations, all seizure clusters (21 clusters across all TeTX animals) started with seizures which then converted to SD events, and were followed by 2-3 more seizures and multiple SD events per seizure (total 124 SDs for 56 total seizures). The SDs appeared to bridge between and initiate the subsequent seizures. We hypothesize that the prolonged impairment of the extracellular dynamics due to SD therefore forms a feedback mechanism (Gorji and Speckmann, 2004; Vinogradova et al., 2006) that may be implicated in linking seizures and SDs within seizure clusters.

There are many peri and post-ictal phenomena whose underlying mechanisms may be linked to seizure-associated spreading depression, including post-ictal generalized EEG suppression (PGES), postictal amnesia, loss of consciousness (Butler and Zeman, 2015), and death. PGES has been extensively investigated and linked to post-ictal cardio-respiratory dysregulation and impaired cognition (Bateman et al., 2008; Lhatoo et al., 2010; Seyal et al., 2011, 2012). There are hypotheses that SD might be the underlying phenomenon of PGES (Rajakulendran and Nashef, 2015). In the epileptic rats discussed here, SD was confined to hippocampus. Nevertheless, postictal SD coincided with prolonged suppression of both hippocampal and cortical activity (Fig. 8A).

Sudden unexplained death in epilepsy (SUDEP) affects approximately 1 in 1000 adults with epilepsy per year (Thurman et al., 2014). Recent studies by Aiba et al. (Aiba and Noebels, 2015) implicate brainstem SD as a mechanism of SUDEP, and have been supported by recent in-vivo recording and imaging studies (Loonen et al., 2019). Clinically, seizure-associated SD has been proposed as a biomarker of imminent SUDEP (Lhatoo et al., 2015). Although we observed a level of modulation of cardiac dynamics during SD (Fig. 8D), in our experiments none of the epileptic rats (n = 6) presented with SUDEP and we identified only one SUDEP instance (Ssentongo et al., 2017) in post-cerebral malaria epileptic mice (n = 21). Therefore in these models, seizure-associated SD does not appear to be a biomarker for SUDEP.

In epileptic rats we found that the estimated propagation speed of SD across the two hippocampi was much faster than what is observed in in vivo or in vitro experiments. In these experiments SD or seizures are often initiated in otherwise normal or resting brain. Although our faster propagation speed might be because the seizure independently initiates SD in multiple places, the stereotyped space-time progression we observe (similar to Fig. 7A), is more consistent with a single SD initiation zone. We therefore hypothesize that the seizure itself primes the tissue through a combination of increases in extracellular potassium, cell swelling, and decreases in available tissue oxygen to support faster propagation. This is analogous to the way that seizure propagation speeds can be parametrically modulated via neuronal excitability (Pinto and Ermentrout, 2001; Richardson et al., 2005).

Interestingly, in the murine model of post-CM chronic epilepsy we estimated slower SD propagation rates than in the TeTX rat model. First, the electrode coverage in the post-CM epilepsy model was not as expansive as that of the TeTX model. Therefore, it could simply be that we did not catch the SD travel path in the post-CM model as well as we did in the TeTX model. Second, the seizure rate, duration, origin, and evolution are different between the two models. These might all be factors in modulating tissue excitability and seizure-induced susceptibility to SD.

SD events are shown to generate long bouts of acidosis (Tschirgi et al., 1957; Somjen, 1984). It is therefore important to note that our observations are not artifacts of the pH dynamics. The μRC EIROF electrodes are pH sensitive (Papeschi et al., 1976). The expected change from SD acidosis should yield positive deflections in the electrode potential. Therefore, our observations of negative deflections in potential were not pH artifacts, and indeed the true magnitude of the physiological shift in tissue potential due to SD was likely larger than we had measured.

We found that spontaneous seizures frequently initiate SD events (in 6/6 epileptic rats: 322 SD events out of 1221 seizures outside of seizure clusters; and 15/21 epileptic mice: 172 SD events out of 441 seizures). Because we observed seizure-associated SD in two different animal models of acquired chronic epilepsy, we conclude that our observations are not simply due to mechanisms of a particular animal model. The combination of our experimental findings and previous modeling efforts motivate/lay out further investigation to uncover the underlying mechanisms that link SD to seizure susceptibility and epilepsy. These results suggest that epilepsy has a broader continuum than just seizures and that SD is a less observed dynamic of epilepsy. Future experiments should address the role of SD in seizure initiation, termination, and potentially intervention.

## Acknowledgments

We thank Ali Nabi, Balaji Shanmugasundaram, and Myles W. Billard for assistance in development of the acquisition hardware.

## Author contributions

F.B., S.J.S., B.J.G. designed research; P.S., J.K., F.B., C.C., and B.J.G. performed surgeries and animal care; J.L. and B.J.G. designed electronics and hardware; F.B. analyzed data; F.B., S.J.S., and B.J.G. wrote the manuscript.

## Data and materials availability

All data and code associated with figures are available upon request, including hour-long segments of raw EEG from which each trace is generated.

